# Elasticity and Topography-Controlled Collagen Hydrogels Mimicking Native Cellular Milieus

**DOI:** 10.1101/706952

**Authors:** Tomoko Gowa Oyama, Kotaro Oyama, Atsushi Kimura, Fumiya Yoshida, Ryo Ishida, Masashi Yamazaki, Hiromi Miyoshi, Mitsumasa Taguchi

**Affiliations:** Quantum Beam Science Research Directorate, National Institutes for Quantum and Radiological Science and Technology (QST), 1233 Watanukimachi, Takasaki-shi, Gunma 370-1292, Japan; PRESTO, Japan Science and Technology Agency (JST), 4-1-8 Honcho, Kawaguchi-shi, Saitama 332-0012, Japan; Graduate School of Science and Technology, Gunma University, 1-5-1 Tenjin-cho, Kiryu-shi, Gunma 376-0052, Japan; Graduate School of Systems Design, Tokyo Metropolitan University, 1-1 Minami-Osawa, Hachioji-shi, Tokyo 192-0397, Japan

**Keywords:** hydrogels, collagens, extracellular matrix, elasticity, topography

## Abstract

Accumulating evidence demonstrates that the elasticity and topography of a cell culture substrate influence cell behavior, in addition to its chemical composition. However, cellular responses to in vivo extracellular matrix (ECM), a hydrogel of proteins (mainly collagen) with various elasticity and a nanometer-to micrometer-scale topography, remain to be elucidated owing to a lack of substrate that provides such complex cues. This study introduces novel collagen hydrogels that can combine, for the first time, elastic, topographic, and compositional cues that recapitulate native ECM. A simple and reagent-free method based on radiation crosslinking alters ECM-derived collagen solutions into hydrogels with a well-defined and tunable elastic modulus covering the broad range of soft tissues (1–236 kPa) and microtopographies while ensuring intrinsic biological functionality of collagen. These collagen hydrogels enabled investigating cell responses to soft topographic cues such as those encountered in vivo, revealing that topography overrides the elasticity and structurally constrains cell morphology by controlling actin cytoskeleton organization. The collagen hydrogels not only reduce in vivo and in vitro behavioral disparity of cells by mimicking native ECM but also facilitate the design of artificial ECM to control cell function and fate in tissue engineering and regenerative medicine.

---

Both elasticity and topographic characteristics of a cell culture substrate influence cell morphogenesis, migration, proliferation, apoptosis, and differentiation, in addition to its chemical composition.^[1–5]^ Thus, altering substrate properties is a non-invasive method for manipulating cell function and fate, and provides temporal and spatial stability, in contrast to biochemical control using growth factors and hormones.^[6]^ As such, this has become one of the major strategies for drug development, and precision and regenerative medicine. However, cells cultured on standard substrates such as plastics and glass exhibit behaviors that differ from those observed in vivo, such as decreased differentiation activity of stem cells, induction of tumorigenesis of healthy cells, and aberrant drug responses.^[7–9]^ To reduce this in vivo and in vitro behavioral disparity, it is essential to use a culture substrate that chemically and physically mimics the native extracellular matrix (ECM). The ECM is a water-containing mesh of proteins (mainly collagen) with an elastic modulus (*E*) of ~1 kPa to a few 100 kPa in soft tissue and a nanometer to micrometer-scale topography. Although a collagen hydrogel can be produced via fibril self-assembly through neutralization and heating a collagen solution, achieving higher stiffness (*E* > 1 kPa) requires chemical cross-linkers that alter cytocompatibility and biodegradability.^[8,10]^

The widely used Matrigel (a protein-mixed hydrogel derived from a mouse tumor) resembles native ECM, but its batch-to-batch biochemical variability introduces a significant level of uncertainty to cell-based experiments.^[11]^ In addition to the elasticity, controlling the topography of ECM-derived hydrogels is technically challenging. Thus, cell responses to topographic cues have mainly been explored on rigid materials (~GPa)^[1,12–14]^ such as silicon and plastics that can be micro-/nano-patterned with techniques such as electron beam lithography and hot embossing. Cellular responses to complex cues provided from native ECM, such as specific topographic cues as soft as the ECM of in vivo soft tissue, remain to be elucidated.

Here, we report collagen hydrogels with well-defined and tunable *E* values, encompassing the broad range of in vivo soft tissue and a microtopography, which was created by the elastic and topographical control by radiation molding (ET-RaM) technique that ensures intrinsic biological functionality of collagen (**Figure 1**a and Experimental Section). Both ECM-derived native collagen and hydrolyzed collagen (gelatin and collagen peptide) can be subjected to ET-RaM. The collagen solution is poured into cell culture dishes (Figure 1a, Step 1), and flexible micropatterned polydimethylsiloxane (PDMS) molds are placed in the solution with cover plates (Figure 1a, Step 2). The samples are then irradiated with ^60^Co γ-rays, which are commonly used to sterilize medical products (Figure 1a, Step 3). Although collagen is decomposed by ionizing radiation in a dried state^[15]^ or at low concentration (<1%),^[16]^ at a certain concentration (a few % to several 10s%) in water (solution or physical gel), collagen is predominantly cross-linked through a reaction with hydroxyl radicals generated in water. ^[15,17]^ Water-containing three-dimensional polymeric networks are thus formed without requiring a cross-linking agent. After removing the PDMS molds, irradiated samples are washed in phosphate-buffered saline (PBS) (Figure 1a, Step 4) and incubated with cell culture medium at 37°C. The collagen hydrogel can then be used for cell culture without any reagent coating (Figure 1a, Step 5). Collagen I forms fibrils through neutralization (Step 4) and heating at 37°C (Step 5). For hydrolyzed collagen, Step 4 is performed at 50°C to remove non-cross-linked components that are soluble in cell culture. Irradiation (Step 3) also sterilizes the sample and can be conducted using other ionizing sources such as an electron beam.

**Figure 1.**
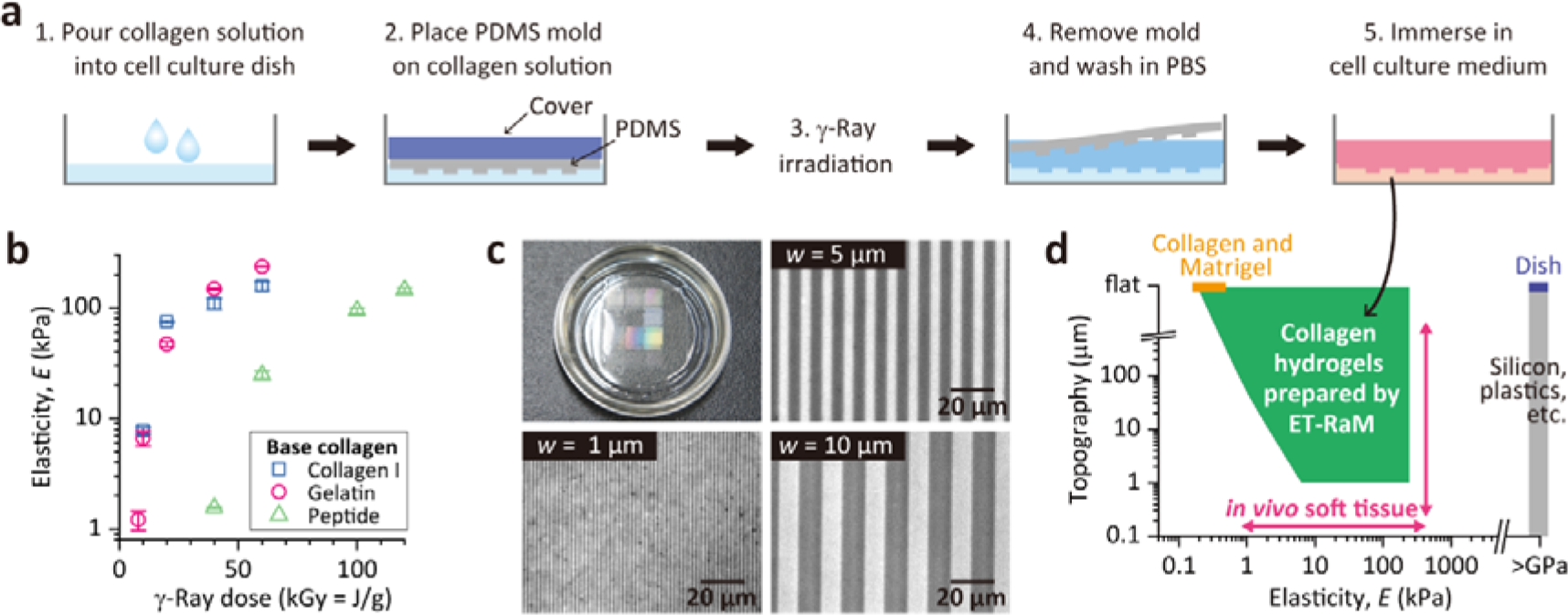
Collagen hydrogels with combined elastic, topographic, and compositional cues that recapitulate native ECM created by ET-RaM without chemical agents. (a) Schematic of ET-RaM process. (b) Controlling *E* of collagen hydrogels by adjusting γ-ray irradiation dose (Step 3). Error bars, SEM (n = 3). (c) Photograph of a typical collagen hydrogel after Step 4 (iridescent reflections are due to microtopography diffraction) and micrographs of representative microtopographies formed on the hydrogel surface. (d) Elasticity and topographic properties of collagen hydrogels compared to conventional cell culture materials^[8,25]^ and in vivo soft tissue.

ET-RaM altered the collagen solutions into hydrogels with regulated stiffness. *E*, determined by the indentation test (Experimental Section and Figure S1a-c, Supporting Information), ranged from 1 to 236 kPa (Figure 1b) without the addition of any reagent. Additionally, *E* showed power-law dependence on the concentration of collagen in hydrogels (Figure S1d, Supporting Information), similar to the reported correlation between in vivo collagen concentration and the *E* of bulk tissue.^[18]^ The volume of collagen I hydrogels was reduced by fibril formation (Steps 4 and 5) (Figure S2a, b, Supporting Information), consistent with the contraction displayed by a fibroblast-containing collagen I hydrogel.^[10,19]^ Collagen I molecules are held in place by cross-linking, forming locally dense cores that promote fibril contraction. The final concentration of 8–157 kPa collagen I hydrogels was 6–14% (Figure 1d, Supporting Information), which cannot be achieved with a conventional collagen I hydrogel (<1%)^[10]^ and is comparable to that in vivo.

With the exception of fluidic hydrogels with *E* ≤ 1 kPa (Figure S2c, Supporting Information), microtopographies were formed on the surface of all types of collagen hydrogels (Figure 1c) with the variation in *E*. Since the above-described compaction of collagen I was isotropic, specific microtopographies can be obtained by adjusting the pattern size of molds (Figure S2a, b, Supporting Information). Mold patterns were directly transferred to gelatin and collagen peptide hydrogels. Thus, our collagen hydrogels covered a broad range of elasticity and topography of native ECM of in vivo soft tissue (Figure 1d).

Another distinguishing feature of the collagen hydrogels is preservation of the biological functionality of collagen, given the essentially unaltered peptide bonds and helical structure (Figure S3a, Supporting Information) while maintaining enzyme-mediated degradability (Figure S3b, Supporting Information). These results were supported by the observation that the amino acid composition ratio was unaffected by ET-RaM (Table S1, Supporting Information). In contrast, chemical cross-linkers specifically consume arginine (R), glutamic acid (E), and aspartic acid (D)^[20,21]^ of cell-binding motifs such as GxxGER and RGD. ET-RaM preserves these motifs and can be applied to various types of collagen—including animal-free recombinant forms—irrespective of the amino acid sequence.

The developed collagen hydrogels can chemically and physically mimic native ECM. Therefore, the collagen hydrogels were used to investigate cell responses to soft topographic cues such as those encountered in vivo. Gelatin was used as the base polymer owing to the broad range of *E* and precise microtopography without the random fibrous structures of collagen I that could affect the cellular response. We analyzed the morphology and actin cytoskeleton organization of Madin-Darby canine kidney (MDCK) epithelial cells, HeLa human cervical carcinoma epithelial cells, and 3T3-Swiss albino mouse embryonic fibroblasts grown on uncoated 7-, 47-, 148-, and 236-kPa collagen hydrogels produced by ET-RaM with a flat surface or with microgrooves (width and spacing = 5 μm, height = 2 μm). On flat hydrogels, the morphology of MDCK, HeLa, and 3T3-Swiss cells changed drastically on day 1 after seeding (Figures S4–S6, Supporting Information). Cells became rounded and formed aggregates over 3 days on the softest (7 kPa) hydrogel. However, with increasing *E*, these cells gradually flattened and spread over a larger area (**Figure 2**a, b and Figures S4–S6, Supporting Information). The flattened cells as well as aggregates adhered to the hydrogels. Proliferation over 3 days was independent of *E* and comparable to that of cells grown in a polystyrene cell culture dish (Figure S7, Supporting Information). MDCK and HeLa cell aggregates partially infiltrated the 7-kPa hydrogels (Figure 2a and Figure S5a, Supporting Information), and 3T3-Swiss cells extended pseudopodia and actively migrated into these hydrogels (Figures S6a and S8, Supporting Information), which is similar to their movement between micropillars.^[22,23]^

**Figure 2.**
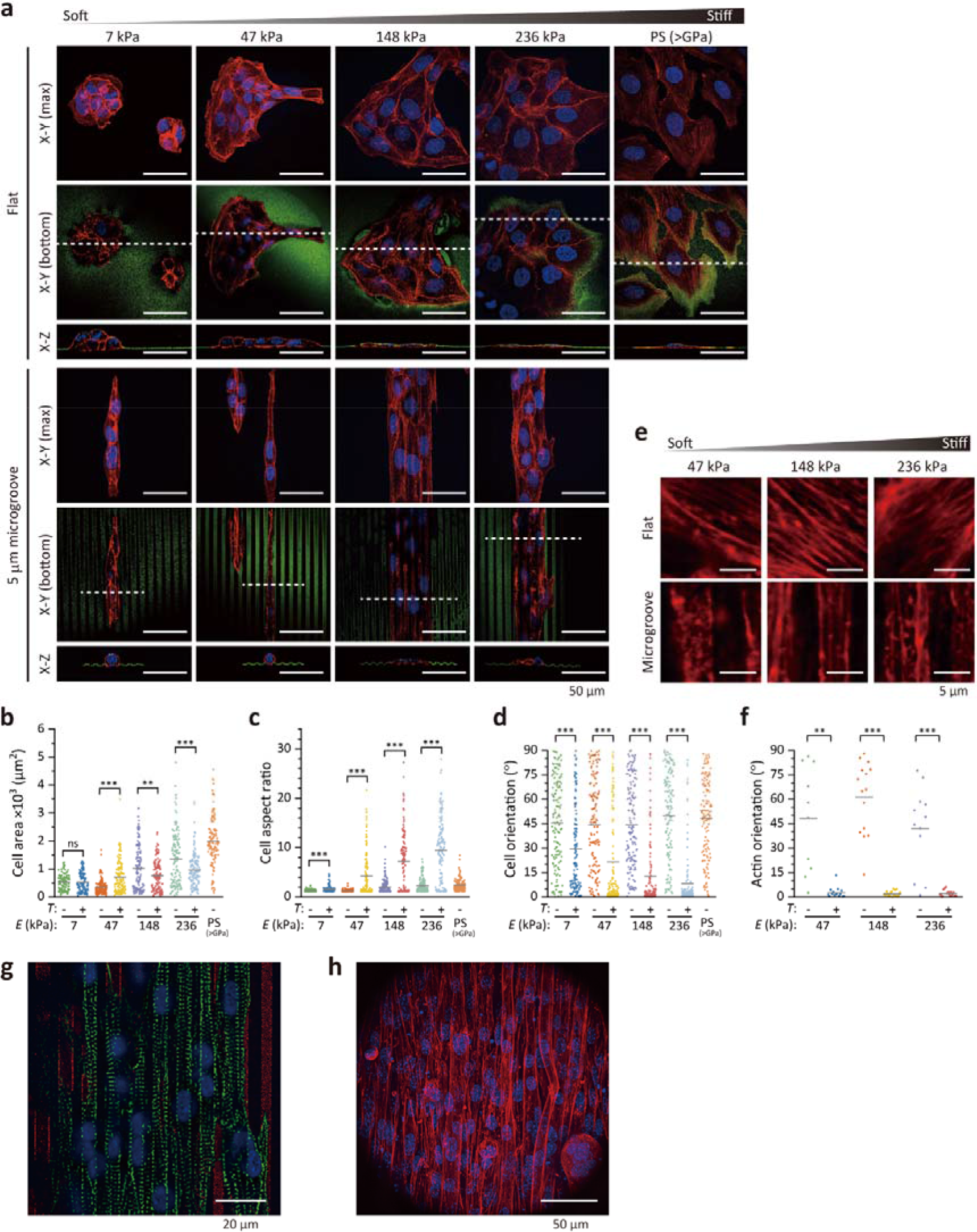
Responses of cells grown on collagen hydrogels prepared by ET-RaM to ECM-mimicking cues. (a) Confocal fluorescence images of nuclear (blue) and actin (red) staining in MDCK cells grown on hydrogels with a flat surface or with 5-μm microgrooves and on a conventional polystyrene (PS) cell culture dish. The hydrogel and PS surfaces were visualized using fluorescent microspheres (green). (b–d) Cell area (b), morphological aspect ratio (c), and orientation (d) of MDCK cells on hydrogels with various elasticity (*E*) values and a flat surface (*T*: −) or 5-μm microgrooves (*T*: +) and in a PS dish are shown. Inserted bars in the graphs indicate the mean values. (e) Actin cytoskeleton beneath the nucleus (red) and (f) actin orientation. Inserted bars in the graphs indicate the mean values. Cells on 7-kPa hydrogels were excluded from analyses due to their multi-layer complexity. Statistical significance in (b–d, f) was assessed with Welch’s t test. ns, not significant (P ≥ 0.05); **P < 0.01, ***P < 0.001 (Table S2, Supporting Information). (g, h) Cardiomyocytes (g) and C2C12 myotubes (h) grown on collagen hydrogels prepared by ET-RaM (236 kPa, 5-μm microgrooves) with nuclear (blue) and sarcomeric α-actinin (green) (g) and nuclei (blue) and actin (red) (h) staining.

Interestingly, the soft topographic cues mimicking the ECM of in vivo soft tissue induced all cell types to spread and become aligned parallel to the microgrooves, even on soft hydrogels where cells formed aggregates on the flat surface (Figure 2a and Figures S4–S6, Supporting Information), except for some 3T3-Swiss cells that migrated into the 7-kPa hydrogels. The morphological aspect ratio and orientation angle relative to the longitudinal direction of the microgrooves differed from those of cells on flat hydrogels on day 1 (Figure 2c, d and Figures S4–S6, Supporting Information); i.e., the cell morphology conformed to microtopographic cues irrespective of *E*. Stiffer hydrogels structurally constrained the alignment of MDCK and 3T3-Swiss cells more effectively owing to the higher aspect ratio and smaller variation in orientation angle. The organization of the actin cytoskeleton was also affected by topographic cues, with actin filaments aligning parallel to the microgrooves (Figure 2e, f and Figures S4–S6, Supporting Information). The collagen hydrogels prepared by ET-RaM were also used to evaluate the response of muscle cells to ECM-mimicking elastic and topographic cues. Rat cardiomyocytes and C2C12 mouse myoblasts seeded directly onto the hydrogels without reagent coating became aligned parallel to the microgrooves (Figure 2g, h), with sarcomeres (the actin-based internal structure) exhibiting a similar orientation (Figure 2g). These cells reacting to soft topographic cues recapitulate their morphology and cytoskeleton observed in vivo.

In summary, we developed collagen hydrogels that can combine elastic, topographic, and compositional cues that recapitulate native ECM, using a simple yet effective method suitable for mass production. Using these collagen hydrogels, we revealed that soft topographic cues, similar to those derived from in vivo ECM, affected cell morphology and actin cytoskeleton organization. These collagen hydrogels are anticipated to reduce in vivo and in vitro behavioral disparity of cells. Moreover, given that elasticity and topographic characteristics are two major cues modulating cell function and fate in tissue formation, maintenance, regeneration, and repair,^[1–5]^ our collagen hydrogels can facilitate the design of artificial ECM to control cell behavior in tissue engineering and regenerative medicine.

## Experimental Section

### ET-RaM of collagen

PDMS molds were prepared in advance as follows. The precursor of PDMS (SIM-260) was mixed with a curing agent (CAT-260) (both from Shin-Etsu Chemical) at a 10:1 ratio, and then degassed and spin-coated on a silicon master mold (DTM1-1; Kyodo International) at 1000 rpm. After curing at 150°C for 30 min, the molds were peeled from the master mold. We used collagen I (porcine skin, 5 mg ml^−1^ [pH 3] solution; Collagen BM, #639-30861; Nitta Gelatin), gelatin (porcine skin, Type A, G1890; Sigma-Aldrich), and peptide (porcine skin, Type A; Nitta Gelatin). The collagen I solution was concentrated to 50 mg ml^−1^ by evaporating the solvent at room temperature (~25°C). The gelatin and peptide solutions were prepared by dissolving in deionized water (supplied by a Millipore Milli-Q system) at 10 wt % by heating at 50°C for 30 min. After pouring the solution into cell culture dishes (Iwaki 1000-035, non-treated, 35 mm; AGC Techno Glass) (Figure 1a, Step 1), micro-patterned flexible PDMS molds loaded onto a plastic cover such as polyethylene-terephthalate film were placed in the solution (Figure 1a, Step 2). The samples were stored overnight at 4°C for collagen I and peptide and at 20°C for gelatin. During storage, gelatin and peptide underwent physical gelation. The samples in sealed bags were irradiated with ^60^Co γ-ray (^60^Co No. 2 Irradiation Facility of Takasaki Advanced Radiation Research Institute, QST) in air at 15–20°C at a dose rate of 10 kGy h^−1^ (kGy = J g^−1^) for ≥ 10-kGy and at 8 kGy h^−1^ for 8-kGy irradiation (Figure 1a, Step 3). The samples were detached from the PDMS molds and immersed in PBS (pH 7.4, 10010023; Thermo Fisher Scientific) (Figure 1a, Step 4) at 37°C for ≥ 3 days to allow collagen I to undergo fibril formation and reach an equilibrium phase, or at 50°C for 2–3 h to remove non-cross-linked components of gelatin and peptide. Collagen hydrogels prepared by ET-RaM were used for cell culture after immersion in culture medium at 37°C for 1 h to replace the absorbed PBS with medium (Figure 1a, Step 5).

### Characterization of collagen hydrogels prepared by ET-RaM

To determine the collagen concentration in hydrogels, swollen samples after Step 4 were filtered through a stainless steel 200-mesh filter (mesh size: 75 μm) and weighed (*W*_s_). After vacuum-drying overnight at 30°C and weighing the dried sample (*W*_g_), the collagen concentration (%) was calculated as (*W*_g_/*W*_s_) × 100.

The elasticity of collagen hydrogels was measured using an indentation tester (RE2-3305B; Yamaden) at 37°C following Step 5. The samples were compressed to 30% deformation with a 2-N load cell using a plunger (φ3 mm or φ8 mm) at 50 μm s^−1^. The compressive modulus was then determined from the slope of stress-strain curves over the strain range from 0% (surface) to the % value equivalent to a depth of 100 μm in each sample.

Photographs of hydrogels and bright-field images of hydrogel microtopography (Figure 1c and Figure S2, Supporting Information) were obtained using a digital camera (DMC-TZ85; Panasonic) and an inverted microscope (IX83; Olympus) with a digital camera (AdvanVision), respectively.

The chemical structures of gelatin before and after the ET-RaM process were compared by Fourier transform infrared (FT-IR) spectroscopy. As the pristine sample, 10 wt% gelatin physical gel was vacuum-dried overnight at 30°C. The hydrogels produced by ET-RaM process were washed in Milli-Q water at 50°C for 2–3 h and vacuum-dried overnight at 30°C. FT-IR spectra were collected using an IRAffinity-1S instrument (Shimadzu) with a DuraSampl IR-II single-reflection diamond attenuated total reflection attachment (Smiths Detection) at a resolution of 1 cm^−1^, and averaged over 32 scans.

The biodegradation properties of gelatin hydrogels were analyzed using proteinase from *Aspergillus oryzae* P0538 (Tokyo Chemical Industry). The pristine sample and hydrogels produced by ET-RaM after washing in Milli-Q water at 50°C for 2–3 h were cut into 10 × 20 × 5-mm pieces and vacuum-dried overnight at 30°C. The samples were weighed (*W*_0_) and then incubated in 0.02 _M_ NaHCO_3_ (#191-01305)/0.001 M CaCl_2_•2H_2_O (#031-00435) (both from Fujifilm Wako Pure Chemical) buffer solution with 50 μg ml^−1^ proteinase in 35-mm dishes at 37°C; they were then filtered through a stainless steel 200-mesh filter and washed with Milli-Q water. After vacuum-drying overnight at 30°C, the samples were weighed (*W*_t_) and % degradation was calculated as [(*W*_0_ − *W*_t_)/*W*_0_] × 100.

Amino acid analysis of gelatin hydrogels was carried out by acidic hydrolysis followed by high-performance liquid chromatography (HPLC). After irradiation in Step 3, samples were washed in Milli-Q water at 50°C for 48 h and vacuum-dried at 30°C for 24 h. The samples and pristine gelatin powder (1 mg each) in separate 1.5-ml glass test tubes were transferred to a 30-ml reaction vial containing 1 ml of 6 M HCl (080-01066; Fujifilm Wako Pure Chemical). The reaction vial was tightly closed in a nitrogen-saturated atmosphere and heated at 110°C for 24 h. The test tubes were then vacuum-dried to remove any remaining HCl, and the obtained hydrolysates were filtered through a 0.45-μm membrane (SFCA033045S; AS ONE) after adding 1 ml of Milli-Q water to each tube. Amino acid standard samples were prepared from an amino acid mixture standard solution (013-08391; Fujifilm Wako Pure Chemical) containing 2.5 × 10^−6^ M of 17 amino acids diluted 20 times with Milli-Q water and filtered through a 45-μm membrane. Borate buffer (pH 9.50) was prepared by mixing 0.05 M boric acid (021-02195) and 0.05 M NaOH (198-13765) (both from Fujifilm Wako Pure Chemical) at a 2:1 volume ratio. Each 10-μl sample solution was mixed with 20 μl of borate buffer and 20 μl of 0.05 M 4-fluoro-7-nitrobenzofurazan (excitation/emission: 470/530 nm, 342-04751; Dojindo Laboratories)/acetonitrile (015-08633; Fujifilm Wako Pure Chemical) and centrifuged at a low speed. The solutions were heated at 60°C for 1 min and then cooled at −18°C for 2 min; 50 μl of 0.3 M HCl was added to terminate the labeling reaction. A 10-μl volume of labeled sample was analyzed by HPLC (Model 1100; Agilent Technologies) with an InertSustainSwift C18 column (GL Sciences) and fluorescence detector (2475; Waters) at 40°C. The HPLC column was eluted with 0.1% trifluoroacetic acid (208-02741; Fujifilm Wako Pure Chemical) and acetonitrile with the gradient mixture ratio recommended by GL Sciences (data no. LB316-0848) at a flow rate of 1.0 ml min^−1^.

### Cell culture on collagen hydrogels prepared by ET-RaM

HeLa (RCB0007), MDCK (RCB0995), and 3T3-Swiss albino mouse embryonic fibroblasts (RCB1642) were obtained from RIKEN BRC Cell Bank. C2C12 cells (EC91031101) were obtained from DS Pharma Biomedical. Cardiomyocytes were obtained from neonatal Wistar rats (Japan SLC) according to our published protocol.^[24]^ Animal experiments were approved by the Institutional Animal Care and Use Committee of QST (approval no. 18-T001-1), and were performed in accordance with the Fundamental Guidelines for Proper Conduct of Animal Experiments and Related Activities in Academic Research Institutions under the jurisdiction of the Ministry of Education, Culture, Sports, Science and Technology of Japan.

HeLa and MDCK cells were cultured in Minimum Essential Eagle’s Medium (MEM) (M5650; Sigma-Aldrich) supplemented with 10% fetal bovine serum (FBS) (SH30910.03; GE Healthcare Japan and 10437-028; Thermo Fisher Scientific, respectively), 100 U ml^−1^ penicillin, 100 μg ml^−1^ streptomycin (15140-122; Thermo Fisher Scientific), and 0.002 M L-glutamine (10378016; Thermo Fisher Scientific and G7513; Sigma-Aldrich, respectively). 3T3-Swiss cells, C2C12 cells, and cardiomyocytes were cultured in Dulbecco’s modified Eagle’s medium (DMEM, 08488-55; Nacalai Tesque) supplemented with 10% FBS, 100 U ml^−1^ penicillin, 100 μg ml^−1^ streptomycin, and 0.002 M L-glutamine (Sigma-Aldrich). A 2-ml volume of each cell suspension (1 × 10^4^ HeLa, MDCK, or 3T3-Swiss cells ml^−1^), 5 × 10^4^ cells ml^−1^ (C2C12), or 1.5 × 10^5^ cells ml^−1^ (cardiomyocytes) prepared using the Countess II FL Automated Cell Counter (AMQAF1000; Thermo Fisher Scientific) was introduced onto collagen hydrogels in 35-mm cell culture dishes without any reagent coating. To induce the differentiation of C2C12 cells, the culture medium was replaced with differentiation medium composed of DMEM supplemented with 2% horse serum (26050-070; Thermo Fisher Scientific), 100 U ml^−1^ penicillin, 100 μg ml^−1^ streptomycin, 0.002 M L-glutamine (Sigma-Aldrich), and 1 μg ml^−1^ insulin (I0516; Sigma-Aldrich) 2 days after seeding. The differentiation medium was refreshed every 1–2 days. Cells were cultured at 37°C in 5% CO_2_.

### Fluorescence staining and imaging

Nuclei, the actin cytoskeleton, and sarcomeres were fluorescently labeled to analyze cell responses to collagen hydrogels. On Day 3 (HeLa, MDCK, and 3T3-Swiss), 6 (C2C12), or 4 (cardiomyocytes) of cell culture, samples were washed with PBS, and cells were fixed with 4% paraformaldehyde (163-20145; Fujifilm Wako Pure Chemical) for 10–15 min. After washing with PBS, the cells were incubated in PBS containing 0.1% Triton X-100 (35501-02; Nacalai Tesque) for 5 min and washed with PBS. HeLa, MDCK, and 3T3-Swiss cells were incubated in PBS containing 0.2 μg ml^−1^ tetramethylrhodamine B isothiocyanate (TRITC)-conjugated phalloidin (P1951; Sigma-Aldrich), 2 μg ml^−1^ 4’,6-diamidino-2-phenylindole (DAPI) (D523; Dojindo Laboratories), and 1% bovine serum albumin (BSA) (019-27051; Fujifilm Wako Pure Chemical) for 20 min. C2C12 cells were incubated in PBS containing 0.1 μg ml^−1^ TRITC-conjugated phalloidin, 1 μg ml^−1^ DAPI, and 1% BSA (P-6154; Biowest) for 20 min. To visualize sarcomeres, cardiomyocytes were incubated in blocking solution composed of PBS with 1% BSA (Fujifilm Wako Pure Chemical) for 60 min, followed by blocking solution containing mouse anti-α-actinin antibody (1:300, A7811; Sigma-Aldrich) for 60 min. After washing with blocking solution, cardiomyocytes were incubated in blocking solution containing Alexa Fluor Plus 488-conjugated goat anti-mouse IgG (1:300, A32723; Thermo Fisher Scientific) for 60 min. To visualize the surface of collagen hydrogels, samples were washed with PBS and incubated for 10 min in PBS containing 1 mg ml^−1^ of 0.2 μm fluorescent microspheres (F8811; Thermo Fisher Scientific). The samples were then washed with PBS and examined with an upright microscope (BX61WI) with a disk scanning unit (BX-DSU), objective lens (LUMPLFLN 60XW) (all from Olympus), and a complementary metal oxide semiconductor camera (ORCA-Flash4.0 V3; Hamamatsu Photonics K.K.). Staining and observation were performed at room temperature (~25°C). Three-dimensional (3D) image stacks were deconvoluted using cellSens (Olympus). Image processing and 3D rendering were performed using ImageJ software (National Institutes of Health).

### Analysis of cell morphology and actin cytoskeleton orientation

Cell morphology was analyzed in terms of spread area, aspect ratio (length of major axis/length of minor axis), and orientation (angle from 0° to 90°) from bright-field images randomly acquired on an inverted microscope (IX70; Olympus) with a CellPad E digital camera system (Allied Scientific Pro) using ImageJ software (n = 3). For the proliferation assay, HeLa cells cultured for 2 h or 1–3 days were stained with 1 μg ml^−1^ Hoechst 33342 solution (H342; Dojindo Laboratories). Nuclei were visualized using a light source (130 W mercury lamp, U-HGLGPS) and mirror units (U-MWU2) (both from Olympus), and the number of HeLa cells was counted from randomly acquired fluorescence images (n = 3). The viability of HeLa cells on Day 3 was analyzed after staining with 1 μg ml^−1^ calcein-AM (C396; Dojindo Laboratories) and 1 μg ml^−1^ DAPI. Viable and dead cells were visualized using mirror units (U-MWU2 and NIBA; Olympus), and percent viability was calculated (n = 3).

The orientation of the actin cytoskeleton was analyzed with a Fourier transformation-based method. An optical section of the actin cytoskeleton beneath the nucleus was first extracted from z-stack images and cropped to 128 × 128 pixels (13.9 μm^2^). The 2D Fourier spectrum for the cropped image was calculated. After removing the direct current offset, a histogram in polar coordinates was generated from the Fourier spectrum data, which was approximated by an ellipse. The orthogonal direction of the major axis of this ellipse was defined as the orientation direction of actin filaments. The angular difference between this and the major axis of microgrooves on collagen hydrogels was defined as the orientation angle of the actin cytoskeleton relative to the microgrooves. The 2D Fourier spectrum was calculated with MATLAB (MathWorks) software, and other image analyses were performed using a macro in ImageJ software.

### Statistics and reproducibility

Data were analyzed with Welch’s t test and one-way analysis of variance followed by a post-hoc Tukey’s honestly significant difference test using OriginPro2018J software (OriginLab). The analyzed datasets of cell responses are summarized in Table S2 (Supporting Information). All experiments were performed at least three times.

## Acknowledgements

The authors thank Ms. Ryoko Mezaki (QST), Ms. Noriko Uchida (QST), and Ms. Bin Jeremiah Duenas Barba (Philippine Nuclear Research Institute) for technical assistance. T.G.O. was supported by Japan Society for the Promotion of Science KAKENHI grants (nos. JP26790070 and JP18K18390). K.O. was supported by a PRESTO grant (no. JPMJPR17P3) from JST. H.M. was supported by an AMED PRIME grant (no. JP18gm5810012). We would like to thank Editage for English language editing.

## References

[1] H. Miyoshi, T. Adachi, Tissue Eng. Part B Rev. 2014, 20, 609.

[2] W. L. Murphy, T. C. McDevitt, A. J. Engler, Nat. Mater. 2014, 13, 547.

[3] Y. Yang, K. Wang, X. Gu, K. W. Leong, Engineering 2017, 3, 36.

[4] K. H. Vining, D. J. Mooney, Nat. Rev. Mol. Cell Biol. 2017, 18, 728.

[5] K. Anselme, N. T. Wakhloo, P. Rougerie, L. Pieuchot, Adv. Healthc. Mater. 2018, 7, 1.

[6] S. W. Cranford, J. De Boer, C. Van Blitterswijk, M. J. Buehler, Adv. Mater. 2013, 25, 802.

[7] C. Yang, M. W. Tibbitt, L. Basta, K. S. Anseth, Nat. Mater. 2014, 13, 645.

[8] S. R. Caliari, J. A. Burdick, Nat. Methods 2016, 13, 405.

[9] G. Brusatin, T. Panciera, A. Gandin, A. Citron, S. Piccolo, Nat. Mater. 2018, 17, 1063.

[10] C. B. Raub, A. J. Putnam, B. J. Tromberg, S. C. George, Acta Biomater. 2010, 6, 4657.

[11] C. S. Hughes, L. M. Postovit, G. A. Lajoie, Proteomics 2010, 10, 1886.

[12] M. J. Dalby, N. Gadegaard, R. Tare, A. Andar, M. O. Riehle, P. Herzyk, C. D. W. Wilkinson, R. O. C. Oreffo, Nat. Mater. 2007, 6, 997.

[13] D. M. Le, K. Kulangara, A. F. Adler, K. W. Leong, V. S. Ashby, Adv. Mater. 2011, 23, 3278.

[14] O. Hasturk, A. Sivas, B. Karasozen, U. Demirci, N. Hasirci, V. Hasirci, Adv. Healthc. Mater. 2016, 5, 2972.

[15] K. Haema, T. G. Oyama, A. Kimura, M. Taguchi, Radiat. Phys. Chem. 2014, 103, 126.

[16] J. P. Miller, B. H. Borde, F. Bordeleau, M. R. Zanotelli, D. J. LaValley, D. J. Parker, L. J. Bonassar, S. C. Pannullo, C. A. Reinhart-King, APL Bioeng. 2018, 2, 031901.

[17] Y. Tomoda, M. Tsuda, Nature 1961, 190, 905.

[18] J. Swift, I. L. Ivanovska, A. Buxboim, T. Harada, P. C. D. P. Dingal, J. Pinter, J. D. Pajerowski, K. R. Spinler, J. W. Shin, M. Tewari, F. Rehfeldt, D. W. Speicher, D. E. Discher, Science 2013, 341.

[19] B. D. Walters, J. P. Stegemann, Acta Biomater. 2014, 10, 1488.

[20] H. W. Sung, D. M. Huang, W. H. Chang, R. N. Huang, J. C. Hsu, J. Biomed. Mater. Res. Part A 1999, 46, 520.

[21] A. Jayakrishnan, S. R. Jameela, Biomaterials 1996, 17, 471.

[22] J. Patterson, J. A. Hubbell, Biomaterials 2010, 31, 7836.

[23] H. Miyoshi, M. Nishimura, Y. Yamagata, H. Liu, Y. Watanabe, M. Sugawara, J. Biomech. Sci. Eng. 2017, 12, 1.

[24] S. A. Shintani, K. Oyama, F. Kobirumaki-Shimozawa, T. Ohki, S. Ishiwata, N. Fukuda, J. Gen. Physiol. 2014, 143, 513.

[25] S. S. Soofi, J. A. Last, S. J. Liliensiek, P. F. Nealey, C. J. Murphy, J. Struct. Biol. 2009, 167, 216.

