## Supporting Information for "Elasticity and Topography-Controlled Collagen Hydrogels Mimicking Native Cellular Milieus"

**Table of contents**

| | |
|---|---|
| Figure S1 | Evaluation of elasticity of collagen hydrogels prepared by ET-RaM and its dependence on concentration. |
| Figure S2 | Limitation of pattern transfer accuracy in the ET-RaM process. |
| Figure S3 | Chemical structures and biodegradation properties of collagen hydrogels prepared by ET-RaM. |
| Figure S4 | Morphology of MDCK cells grown on collagen hydrogels prepared by ET-RaM. |
| Figure S5 | Responses of HeLa cells grown on collagen hydrogels prepared by ET-RaM to ECM-mimicking cues. |
| Figure S6 | Responses of 3T3-Swiss albino cells grown on collagen hydrogels prepared by ET-RaM to ECM-mimicking cues. |
| Figure S7 | Proliferation and viability of HeLa cells grown on collagen hydrogels prepared by ET-RaM. |
| Figure S8 | Migration of 3T3-Swiss albino cells into collagen hydrogels prepared by ET-RaM. |
| Table S1 | Amino acid analysis of collagen hydrogels prepared by ET-RaM |
| Table S2 | Datasets of cell responses analyzed in this study |

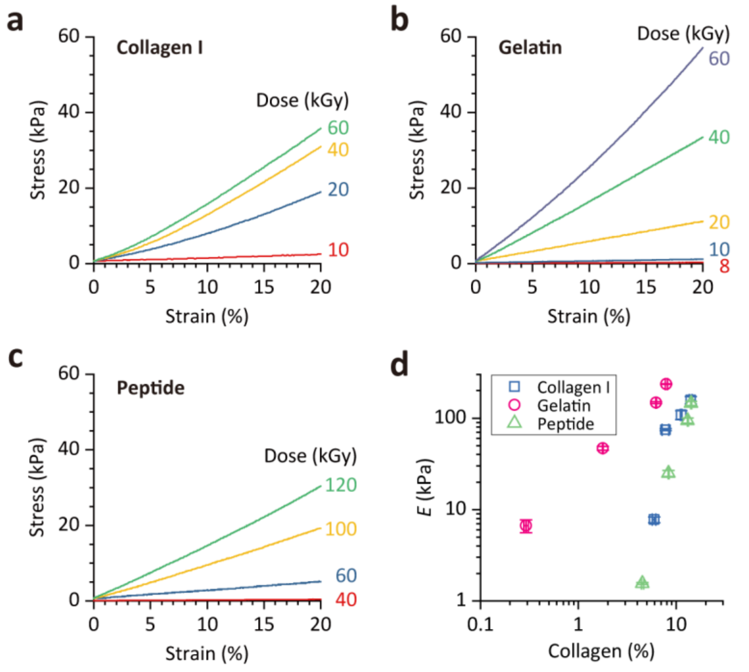


**Figure S1. Evaluation of elasticity of collagen hydrogels prepared by ET-RaM and its dependence on concentration.** (a–c) Representative stress-strain curves of collagen hydrogels obtained with collagen I (a), gelatin (b), and collagen peptide (c) as a base polymer. Hydrogel elasticity (*E*) in terms of elastic modulus was determined from the slope of the approximately straight lines of the stress-strain curve over a strain ranging from 0% (surface) to the % value equivalent to a depth of 100 μm in each sample. (d) *E* values showing power-law dependence on collagen concentration in collagen hydrogels, except for the softest hydrogel obtained with collagen I. Error bars for *E* and collagen concentration represent SEM (n = 3).

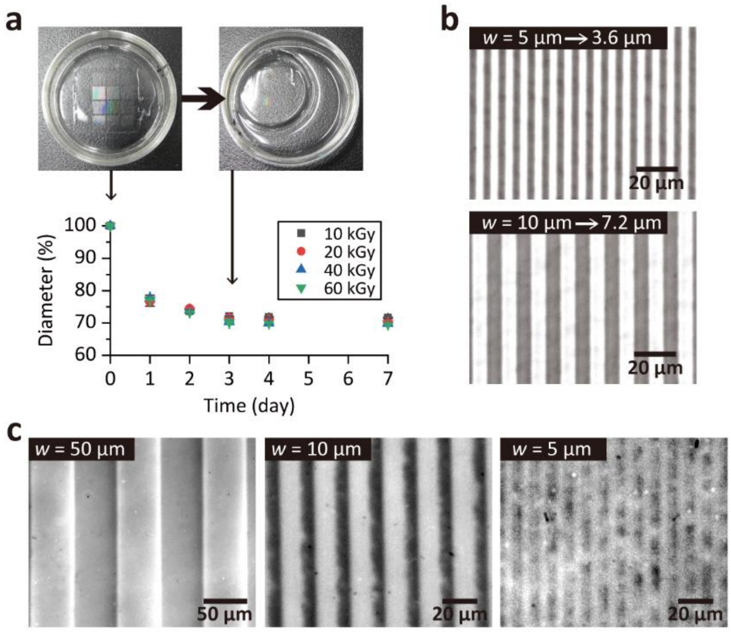


**Figure S2. Limitation of pattern transfer accuracy in the ET-RaM process.** (a) Photograph of a typical collagen I hydrogel prepared by ET-RaM irradiated at 20 kGy $^{60}$Co γ-ray before and after neutralization (Step 4) and heating at 37°C (Step 5) of the ET-RaM process. Compaction was determined from the diameter. Error bars represent SEM (n = 3). (b) Micrographs of representative topographies formed on the surface of compacted collagen I hydrogels irradiated at 20 kGy. The width of transferred microgrooves decreased from 5 and 10 µm to 3.6 and 7.2 µm, respectively. (c) Micrographs of representative topographies formed on the collagen hydrogel surface with an *E* of 1.2 kPa (base polymer: gelatin, irradiation dose: 8 kGy). Precise control of microtopography was limited for ≤ 1 kPa fluidic hydrogels.

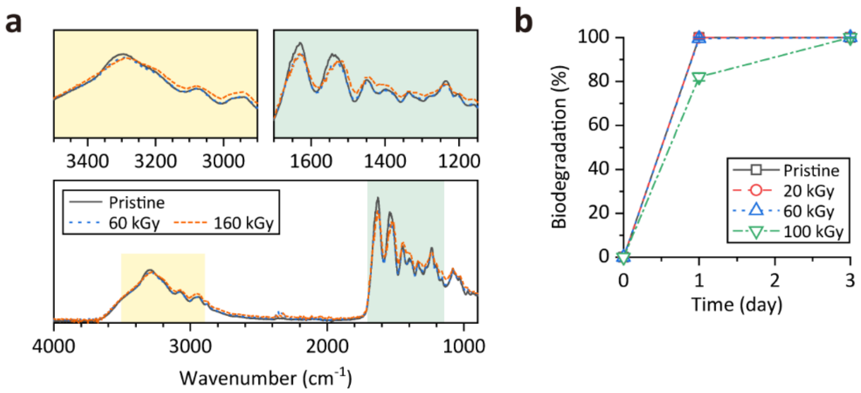


**Figure S3. Chemical structures and biodegradation properties of collagen hydrogels prepared by ET-RaM.** (a) Fourier-transform infrared spectra of gelatin before and after ET-RaM processing with irradiation doses of 60 and 160 kGy. The amide bands of polypeptides (A: 3297 $cm^{-1}$, B: 3075 $cm^{-1}$, I: 1630 $cm^{-1}$, II: 1537 $cm^{-1}$, and III: 1237 $cm^{-1}$) related to peptide bonds and helical structure were largely unaltered by ET-RaM. A slight shift to a lower wave number for amide A and amide II suggests a small change in the secondary structure of gelatin. (b) In vitro enzymatic biodegradation of the gelatin hydrogel evaluated using proteinase (n = 3). Biodegradability was maintained after the ET-RaM process although it was slowed down at radiation doses > 100 kGy.

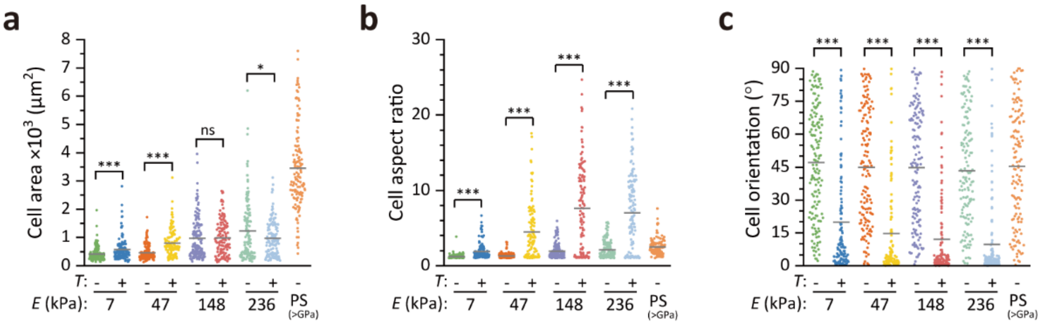


**Figure S4. Morphology of MDCK cells grown on collagen hydrogels prepared by ET-RaM.** (a–c) Cell area (a), morphological aspect ratio (b), and orientation (c) of MDCK cells grown on collagen hydrogels with varying elasticity (*E*) and a flat surface (*T*: –) or 5-µm microgrooves (*T*: +) or grown in a polystyrene (PS) dish on Day 1. Inserted bars indicate the mean values. Statistical significance was assessed with Welch's t test. ns, not significant ($P \geq 0.05$); *$P < 0.05$, ***$P < 0.001$ (Table S2). On Day 1 after seeding, MDCK cells already exhibited altered morphology in response to changes in *E* and microtopography.

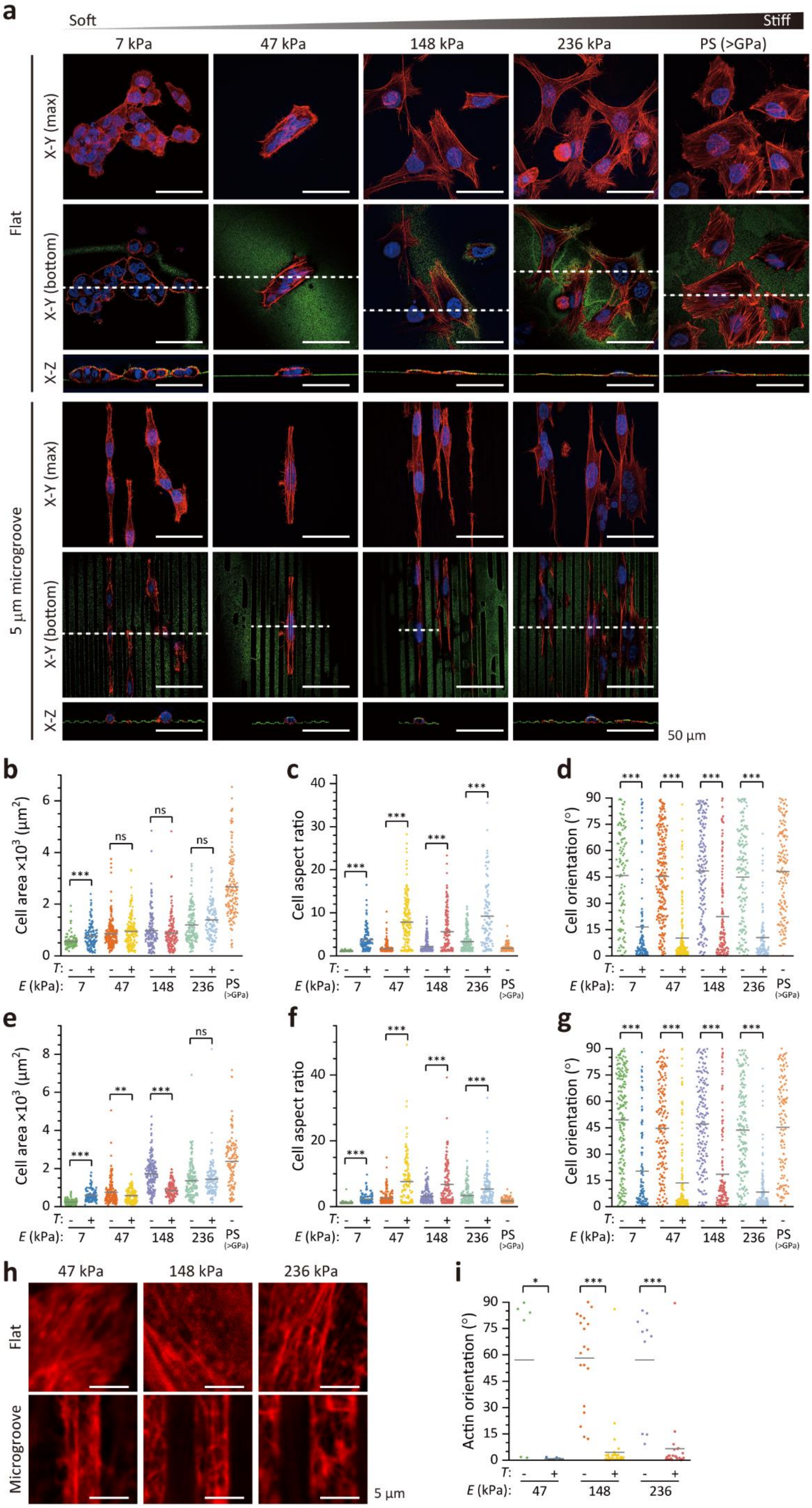
a
Soft
Stiff
7 kPa
47 kPa
148 kPa
236 kPa
PS (>GPa)
Flat
X-Y (max)
X-Y (bottom)
X-Z
5 μm microgroove
50 μm
b
Cell area ×10³ (μm²)
c
Cell aspect ratio
d
Cell orientation (°)
e
Cell area ×10³ (μm²)
f
Cell aspect ratio
g
Cell orientation (°)
T:
E (kPa):
ns
h
47 kPa
148 kPa
236 kPa
Flat
Microgroove
5 μm
i
Actin orientation (°)

**Figure S5. Responses of HeLa cells grown on collagen hydrogels prepared by ET-RaM to ECM-mimicking cues.** (a) Confocal fluorescence micrographs of nuclei (blue) and actin (red) staining in HeLa cells grown on collagen hydrogels with a flat surface or with microgrooves, or grown in a polystyrene (PS) cell culture dish. Hydrogel and PS surfaces were visualized with fluorescent microspheres (green). (b–g) Area, morphological aspect ratio, and orientation of HeLa cells grown on collagen hydrogels with varying elasticity (*E*) and a flat surface (*T*: –) or 5-μm microgrooves (*T*: +), or grown in a PS dish on Day 1 (b–d) and Day 3 (e–g) are shown. Inserted bars in the graphs indicate the mean values. (h, i) Actin cytoskeleton beneath the nucleus (red) (h) and actin orientation angle (i). Inserted bars in the graphs indicate the mean values. Statistical significance in (b–g and i) was assessed with Welch's t test. ns, not significant ($P \geq 0.05$); *$P < 0.05$, **$P < 0.01$, ***$P < 0.001$ (Table S2).

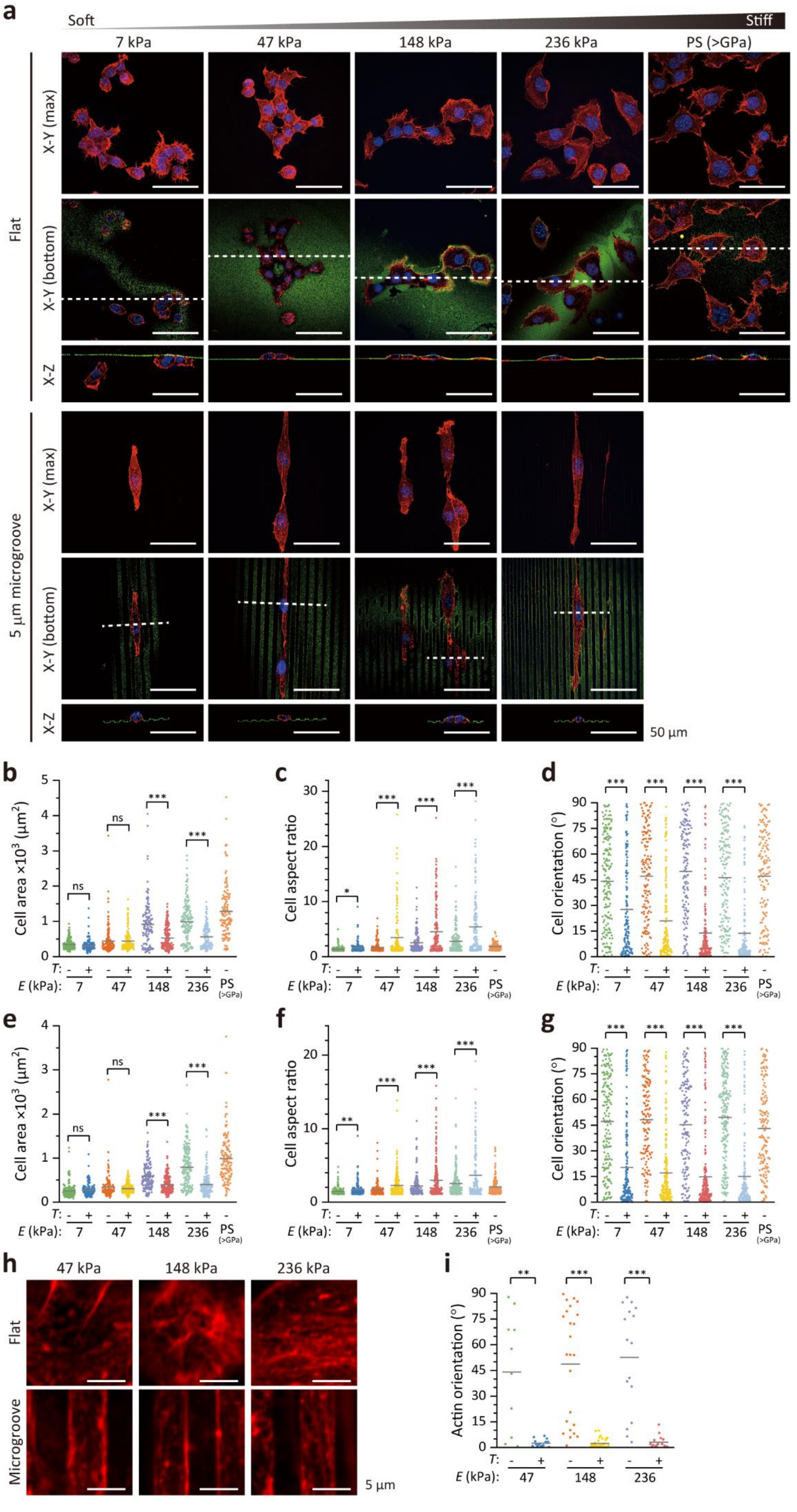
a
Soft
Stiff
7 kPa
47 kPa
148 kPa
236 kPa
PS (>GPa)
Flat
X-Y (max)
X-Y (bottom)
X-Z
5 μm microgroove
50 μm
b
Cell area ×10$^3$ (μm$^2$)
c
Cell aspect ratio
d
Cell orientation (°)
e
f
g
T:
E (kPa):
ns
*
**
***
h
47 kPa
148 kPa
236 kPa
Microgroove
5 μm
i
Actin orientation (°)

**Figure S6. Responses of 3T3-Swiss albino cells grown on collagen hydrogels prepared by ET-RaM to ECM-mimicking cues.** (a) Confocal fluorescence micrographs of nuclei (blue) and actin (red) staining in 3T3-Swiss cells grown on collagen hydrogels with a flat surface or with microgrooves, or grown in a polystyrene (PS) cell culture dish. Hydrogel and PS surfaces were visualized with fluorescent microspheres (green). (b–g) Area, morphological aspect ratio, and orientation of 3T3-Swiss cells grown on collagen hydrogels with varying elasticity (*E*) and a flat surface (*T*: –) or 5-μm microgrooves (*T*: +), or grown in a PS dish on Day 1 (b–d) and Day 3 (e–g) are shown. Inserted bars in the graphs indicate the mean values. (h, i) Actin cytoskeleton beneath the nucleus (red) (h) and actin orientation angle (i). Inserted bars in the graphs indicate the mean values. Statistical significance in (b–g and i) was assessed with Welch's t test. ns, not significant ($P \geq 0.05$); *$P < 0.05$, **$P < 0.01$, ***$P < 0.001$ (Table S2).

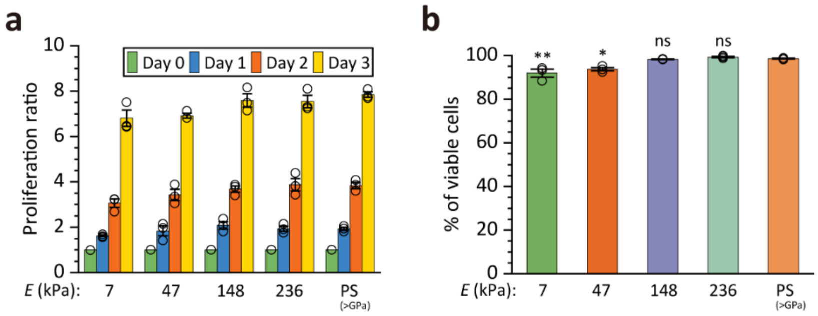


**Figure S7. Proliferation and viability of HeLa cells grown on collagen hydrogels prepared by ET-RaM.** (a) Proliferation over 3 days of HeLa cells grown on a collagen hydrogel (base polymer: gelatin) or a polystyrene (PS) cell culture dish. Error bars represent SEM (n = 3). There were no differences in proliferation among cells grown on collagen hydrogels with different stiffness values and on the PS cell culture dish ($P \geq 0.05$, one-way analysis of variance [ANOVA] followed by a post-hoc Tukey's honestly significant difference [HSD] test) (Table S2). (b) Viability of HeLa cells on Day 3. Error bars represent SEM (n = 3). ns, not significant ($P \geq 0.05$); *$P < 0.05$, **$P < 0.01$ vs. PS cell culture dish (one-way ANOVA followed by a post-hoc Tukey's HSD test) (Table S2).

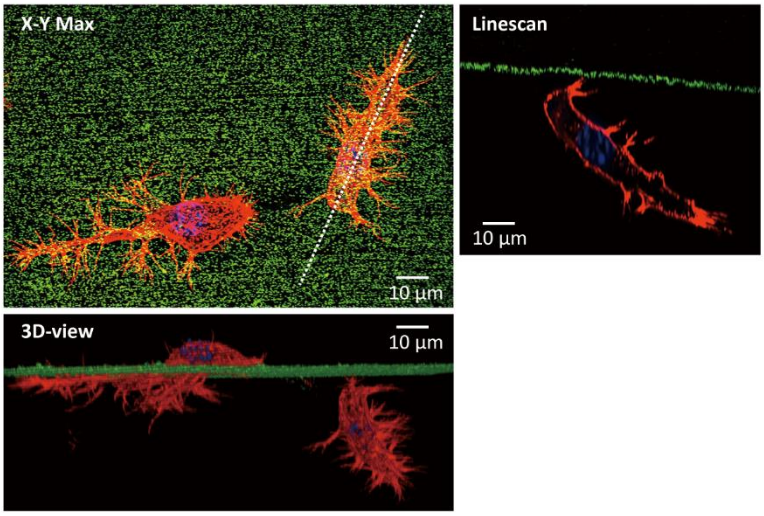


**Figure S8. Migration of 3T3-Swiss albino cells into collagen hydrogels prepared by ET-RaM.** Confocal fluorescence micrographs of nuclear (blue) and actin (red) staining in cells grown on collagen hydrogel with $E$ = 7 kPa.

### Table S1. Amino acids analysis of collagen hydrogels prepared by ET-RaM

| Amino acid | Pristine | ET-RaM with 60 kGy | | ET-RaM with 160 kGy | | |
|---|---|---|---|---|---|---|
| | % | % | P value vs. pristine | % | P value vs. pristine | P value vs. 60 kGy |
| **Glycine** | 33.52 ± 0.66 | 33.88 ± 0.72 | 0.9855 | 32.04 ± 2.56 | 0.7929 | 0.7028 |
| **Proline** | 18.54 ± 1.17 | 19.78 ± 3.83 | 0.9687 | 20.49 ± 4.82 | 0.9241 | 0.9893 |
| **Serine** | 14.60 ± 1.54 | 15.24 ± 1.37 | 0.9502 | 15.94 ± 1.53 | 0.8044 | 0.9408 |
| **Alanine** | 7.83 ± 0.76 | 8.48 ± 1.75 | 0.9564 | 8.25 ± 1.99 | 0.9817 | 0.9943 |
| **Arginine** | 7.72 ± 0.34 | 4.84 ± 2.07 | 0.2890 | 6.60 ± 0.24 | 0.7988 | 0.5904 |
| **Glutamic acid + Glutamine** | 3.86 ± 0.54 | 4.78 ± 1.57 | 0.8617 | 4.23 ± 1.37 | 0.9765 | 0.9462 |
| **Lysine** | 2.77 ± 0.43 | 2.24 ± 0.43 | 0.7782 | 2.19 ± 0.71 | 0.7400 | 0.9974 |
| **Leucine** | 2.70 ± 0.22 | 2.52 ± 0.14 | 0.8102 | 2.36 ± 0.25 | 0.5174 | 0.8606 |
| **Threonine** | 2.16 ± 0.10 | 1.93 ± 0.03 | 0.2508 | 1.86 ± 0.13 | 0.1328 | 0.8723 |
| **Phenylalanine** | 2.12 ± 0.09 | 1.78 ± 0.30 | 0.5270 | 1.55 ± 0.21 | 0.2227 | 0.7469 |
| **Aspartic acid + Asparagine** | 1.45 ± 0.20 | 1.87 ± 0.70 | 0.8481 | 1.65 ± 0.58 | 0.9616 | 0.9559 |
| **Isoleucine** | 1.22 ± 0.10 | 1.12 ± 0.09 | 0.6609 | 1.05 ± 0.02 | 0.3346 | 0.7976 |
| **Histidine** | 1.03 ± 0.24 | 1.16 ± 0.51 | 0.9811 | 1.36 ± 0.67 | 0.8877 | 0.9570 |
| **Tyrosine** | 0.48 ± 0.03 | 0.39 ± 0.05 | 0.6139 | 0.43 ± 0.10 | 0.8424 | 0.9107 |

Composition ratio (%) of 14 amino acids in pristine gelatin hydrogel and gelatin hydrogels prepared by ET-RaM with $^{60}$Co γ-ray irradiation at 60 and 160 kGy. Data represent mean ± SEM (n = 3). There was no significant difference in amino acid composition among pristine and the two types of hydrogel by one-way analysis of variance followed by a post-hoc Tukey's honestly significant difference test.

**Table S2. Datasets of cell responses analyzed in this study.**

Morphological alterations in MDCK cells.

| Day | *E* (kPa) | *T* | No. of cells | Cell area | | Cell aspect ratio | | Cell orientation | |
|---|---|---|---|---|---|---|---|---|---|
| | | | | P value | Related data | P value | Reference | P value | Related data |
| **1** | **7** | – | 150 | 2.00E-4 | Figure S4a | 9.03E-11 | Figure S4b | 1.14E-17 | Figure S4c |
| | | + | 128 | | | | | | |
| | **47** | – | 143 | 1.30E-10 | | 3.31E-15 | | 2.89E-21 | |
| | | + | 121 | | | | | | |
| | **148** | – | 144 | 0.8723 | | 4.93E-20 | | 1.03E-25 | |
| | | + | 129 | | | | | | |
| | **236** | – | 138 | 0.0168 | | 1.40E-20 | | 1.61E-29 | |
| | | + | 135 | | | | | | |
| | **>GPa (PS dish)** | – | 132 | — | | — | | | |
| **3** | **7** | – | 144 | 0.8722 | Figure 2b | 2.43E-5 | Figure 2c | 1.69E-6 | Figure 2d |
| | | + | 128 | | | | | | |
| | **47** | – | 140 | 1.01E-10 | | 2.46E-12 | | 8.57E-11 | |
| | | + | 131 | | | | | | |
| | **148** | – | 144 | 0.0014 | | 1.56E-17 | | 1.46E-23 | |
| | | + | 135 | | | | | | |
| | **236** | – | 138 | 5.27E-5 | | 2.18E-26 | | 1.46E-36 | |
| | | + | 142 | | | | | | |
| | **> GPa (PS dish)** | – | 136 | — | | — | | — | |

PS, polystyrene

Actin filament orientation in MDCK cells

| Day | *E* (kPa) | *T* | No. of cells | P value | Related Data |
|---|---|---|---|---|---|
| **3** | **47** | – | 10 | 0.0015 | Figure 2f |
| | | + | 15 | | |
| | **148** | – | 15 | 6.00E-8 | |
| | | + | 20 | | |
| | **236** | – | 12 | 2.00E-4 | |
| | | + | 20 | | |

### Morphological alterations in HeLa cells

| Day | *E* (kPa) | *T* | No. of cells | Cell area | | Cell aspect ratio | | Cell orientation | |
|---|---|---|---|---|---|---|---|---|---|
| | | | | P value | Related data | P value | Reference | P value | Related data |
| **1** | **7** | – | 107 | 8.80E-5 | Figure S5b | 1.57E-19 | Figure S5c | 9.44E-16 | Figure S5d |
| | | + | 121 | | | | | | |
| | **47** | – | 236 | 0.0646 | | 4.25E-29 | | 1.12E-49 | |
| | | + | 164 | | | | | | |
| | **148** | – | 158 | 0.2424 | | 7.30E-14 | | 3.53E-16 | |
| | | + | 145 | | | | | | |
| | **236** | – | 160 | 0.0223 | | 1.58E-13 | | 7.28E-33 | |
| | | + | 107 | | | | | | |
| | **> GPa (PS dish)** | – | 136 | — | | — | | — | |
| **3** | **7** | – | 193 | 4.67E-20 | Figure S5e | 1.85E-12 | Figure S5f | 4.79E-19 | Figure S5g |
| | | + | 108 | | | | | | |
| | **47** | – | 168 | 0.0023 | | 4.19E-13 | | 2.16E-26 | |
| | | + | 145 | | | | | | |
| | **148** | – | 167 | 4.35E-25 | | 2.63E-9 | | 4.81E-21 | |
| | | + | 136 | | | | | | |
| | **236** | – | 168 | 0.4245 | | 1.56E-5 | | 2.48E-37 | |
| | | + | 138 | | | | | | |
| | **> GPa (PS dish)** | – | 120 | — | | — | | — | |

PS, polystyrene.

### Actin filament orientation in HeLa cells

| Day | *E* (kPa) | *T* | No. of cells | P value | Related data |
|---|---|---|---|---|---|
| **3** | **47** | – | 6 | 0.0241 | Figure S5i |
| | | + | 7 | | |
| | **148** | – | 19 | 1.64E-8 | |
| | | + | 39 | | |
| | **236** | – | 10 | 4.62E-4 | |
| | | + | 24 | | |

### Morphological alterations in 3T3-Swiss albino cells

| Day | *E* (kPa) | *T* | No. of cells | Cell area | | Cell aspect ratio | | Cell orientation | |
|---|---|---|---|---|---|---|---|---|---|
| | | | | **P value** | **Related Data** | **P value** | **Related Data** | **P value** | **Related Data** |
| **1** | **7** | – | 164 | 0.1078 | Figure S6b | 0.0110 | Figure S6c | 2.24E-7 | Figure S6d |
| | | + | 138 | | | | | | |
| | **47** | – | 141 | 0.9126 | | 1.56E-6 | | 3.02E-16 | |
| | | + | 148 | | | | | | |
| | **148** | – | 135 | 1.85E-9 | | 8.68E-7 | | 1.16E-30 | |
| | | + | 167 | | | | | | |
| | **236** | – | 161 | 8.89E-15 | | 5.87E-7 | | 4.63E-27 | |
| | | + | 141 | | | | | | |
| | **> GPa (PS dish)** | – | 137 | — | | — | | — | |
| **3** | **7** | – | 165 | 0.3708 | Figure S6e | 0.0047 | Figure S6f | 8.42E-19 | Figure S6g |
| | | + | 156 | | | | | | |
| | **47** | – | 142 | 0.1597 | | 3.47E-5 | | 1.01E-28 | |
| | | + | 193 | | | | | | |
| | **148** | – | 153 | 1.41E-11 | | 3.95E-4 | | 9.05E-26 | |
| | | + | 210 | | | | | | |
| | **236** | – | 205 | 4.59E-22 | | 1.38E-4 | | 1.13E-36 | |
| | | + | 162 | | | | | | |
| | **> GPa (PS dish)** | – | 151 | — | | — | | — | |

PS, polystyrene.

### Actin filament orientation in 3T3-Swiss albino cells

| Day | *E* (kPa) | *T* | No. of cells | P value | Related Data |
|---|---|---|---|---|---|
| **3** | **47** | – | 10 | 0.0039 | Figure S6i |
| | | + | 15 | | |
| | **148** | – | 24 | 5.04E-7 | |
| | | + | 32 | | |
| | **236** | – | 16 | 1.22E-5 | |
| | | + | 15 | | |

### Proliferation ratio of HeLa cells in Figure S7a

| *E* (kPa) | Data no. | Day 0 | Day 1 | | | Day 2 | | | Day 3 | | |
|---|---|---|---|---|---|---|---|---|---|---|---|
| | | No. of cells | No. of cells | Ratio | P value | No. of cells | Ratio | P value | No. of cells | Ratio | P value |
| **7** | **1** | 124 | 200 | 1.61 | 0.5410 | 334 | 2.69 | 0.1248 | 802 | 6.47 | 0.0882 |
| | **2** | 145 | 228 | 1.57 | | 469 | 3.23 | | 933 | 6.43 | |
| | **3** | 137 | 232 | 1.69 | | 445 | 3.25 | | 1030 | 7.52 | |
| **47** | **1** | 134 | 189 | 1.41 | 0.9865 | 520 | 3.88 | 0.6158 | 912 | 6.81 | 0.1386 |
| | **2** | 130 | 278 | 2.14 | | 434 | 3.34 | | 926 | 7.12 | |
| | **3** | 211 | 414 | 1.96 | | 643 | 3.05 | | 1437 | 6.81 | |
| **148** | **1** | 164 | 388 | 2.37 | 0.9325 | 557 | 3.40 | 0.9785 | 1171 | 7.14 | 0.9450 |
| | **2** | 155 | 279 | 1.80 | | 600 | 3.87 | | 1263 | 8.15 | |
| | **3** | 137 | 285 | 2.08 | | 517 | 3.77 | | 1024 | 7.47 | |
| **236** | **1** | 129 | 245 | 1.90 | 1 | 491 | 3.81 | 1 | 942 | 7.30 | 0.9085 |
| | **2** | 150 | 321 | 2.14 | | 658 | 4.39 | | 1214 | 8.09 | |
| | **3** | 173 | 302 | 1.75 | | 593 | 3.43 | | 1252 | 7.24 | |
| **>GPa (PS dish)** | **1** | 102 | 209 | 2.05 | — | 394 | 3.86 | — | 825 | 8.09 | — |
| | **2** | 117 | 212 | 1.81 | | 420 | 3.59 | | 900 | 7.69 | |
| | **3** | 108 | 209 | 1.94 | | 440 | 4.07 | | 838 | 7.76 | |

PS, polystyrene.

### Viability of HeLa cells in Figure S7b

| *E* (kPa) | Data no. | No. of cells | | Viability (%) | P value |
|---|---|---|---|---|---|
| | | Viable | Dead | | |
| **7** | **1** | 914 | 67 | 93.17 | 0.0022 |
| | **2** | 879 | 58 | 93.81 | |
| | **3** | 1343 | 181 | 88.12 | |
| **47** | **1** | 842 | 58 | 93.56 | 0.0301 |
| | **2** | 913 | 71 | 92.78 | |
| | **3** | 812 | 38 | 95.53 | |
| **148** | **1** | 1262 | 21 | 98.36 | 1 |
| | **2** | 765 | 12 | 98.46 | |
| | **3** | 808 | 13 | 98.42 | |
| **236** | **1** | 527 | 1 | 99.81 | 0.9780 |
| | **2** | 641 | 5 | 99.23 | |

| | | | | | |
|---|---|---|---|---|---|
| | **3** | 562 | 7 | 98.77 | |
| **>GPa (PS dish)** | **1** | 525 | 7 | 98.68 | — |
| | **2** | 516 | 6 | 98.85 | |
| | **3** | 526 | 10 | 98.13 | |

PS, polystyrene.
